# Assessment and phenotypic identification of millet germplasm (*Setaria italica* [L.]) in Liaoning, China

**DOI:** 10.1101/2024.04.14.589429

**Authors:** Li Xin-tong, He Wei-Feng, Wang Hong-hao, Xu Min

**Affiliations:** School of Accounting, GuiZhou University of Finance and Economics, Guiyang 550025, Guizhou, China; Cash Crop Institute, Liaoning Academy of Agricultural Science, Liaoyang 111000, Liaoning, China

**Keywords:** Millet, Germplasm resources, Agronomy trait, Phenotypic identification, Diversity analysis

## Abstract

**Aims:** This study evaluated millet germplasms in Liaoning Province to support the collection, preservation and innovation of millet germplasm resources.

**Methods:** The study was conducted from 2018 to 2020, involved the selection of 105 millet germplasm resources from the Germplasm Bank of the Liaoning Academy of Agricultural Sciences (LAAS), the observation and recording of 31 traits, and the application of multivariate analysis methods to assess phenotypic diversity.

**Results:** From the diversity analysis and correlation analysis, it was found that the tested traits had abundant diversity and complex correlations among them. principal component analysis (PCA) comprehensively analyzed all quantitative traits and extracted 7 principal components. Grey relational analysis (GRA) highlighted the varied contributions of different traits to yield. Through systematic cluster analysis, the resources were categorized into six groups at Euclidean distance of 17.09. K-mean cluster analysis determined the distribution interval and central value of each trait, then identified resources with desirable traits.

**Conclusion:** The results revealed resources that possess characteristics such as upthrow seedling leaves, more tillers and branches, larger and well-formed ears, and lodging resistance prefer to higher grain yield. It was also discovered that the subear internode length (SIL) could be an indicator for maturity selection. Four specific resources, namely, Dungu No. 1, Xiao-li-xiang, Basen Shengu, and Yuhuanggu No. 1, were identified for further breeding and practical applications.

## 1. Introduction

Millet (*Setaria italica* [L.]) is a diploid (2*n*=2*X* =18) species of foxtail grass (*Gramineae, Setaria*). Millet, which originated in China, is one of the world’s oldest cultivated crops, with a cultivation history of 5000 to 8000 years^[1]^. Millet possesses attributes such as drought resistance, water efficiency, high light utilization capacity, high storage convenience, and dual utility as both a grain and a grass^[2]^. The millet seed kernel is a reservoir of well-balanced nutrients^[3]^, comprising ample proteins^[4]^ and vitamins^[5]^. It is commonly chosen as the primary dietary option for new mothers or recovering patients.

According to statistical data, there are more than 40,000 millet resources worldwide. Among these, the country with the richest germplasm resources is China, specifically the National Crop Germplasm Bank, which stores over 26,000 resources, accounting for 70% of the global collection^[4,6]^. Previous researchers have made significant progress in researching and utilizing millet germplasm resources. These studies primarily focused on genetic diversity analysis of phenotypic traits. For instance, Wang^[7]^ comprehensively evaluated 15 phenotypic traits in 878 millet resources globally and identified 8 key indicators, such as leaf sheath color, ear length, seed color, and kernel color, for phenotype identification. Tian^[8]^ investigated the genetic diversity of 482 millet varieties in Henan and Shandong provinces and discovered that the diversity level of millet breeding cultivars was considerably lower than that of local varieties, suggesting certain traits of greater significance in the breeding process. Li^[9]^ examined 23,381 Chinese landrace millet samples and conducted a comprehensive analysis of 11 agronomy traits. Of these traits, only seedling leaf color, starch composition, and 1000-grain weight showed significant regional differences in phenotypic diversity indices. In addition, molecular biotechnology has been applied in the study of millet germplasm resources. For instance, Yang^[10]^ and Schontz^[11]^ observed abundant genetic diversity among millet resources from various regions. Jia^[12]^ conducted genome resequencing to analyse the diversity of 916 core millet germplasms, providing insight into the geographical distribution of millet genetic resources.

According to Liu^[13]^, the primary millet-producing regions are concentrated in the northern and eastern parts of China. The millet produced in Liaoning, an important millet-producing area, is renowned for its golden-red grain color, pleasant taste, and high quality. In comparison to molecular markers, simpler and more intuitive indicators are required to evaluate resource materials in practical production. Therefore, this study aimed to comprehensively understand the basic situation of millet germplasm resources in Liaoning Province, to screen agronomy traits as selection indicators, and to promote millet scientific research and production in northeastern China.

## 2. Materials and Methods

### 2.1 Experimental materials

A total of 105 millet resources preserved in the Germplasm Bank of the Liaoning Academy of Agricultural Sciences (LAAS) were selected for this study. These resources originated from various regions, including Liaoning (52), Beijing (20), Hebei (8), Jilin (9), Inner Mongolia (9), Heilongjiang (2), and Shanxi (5). Approximately half of the materials are from Liaoning, with the remaining resources sourced from provinces that north of the Yellow River. Fig. 1 illustrates the origins of the resources utilized in this study.

**Fig. 1:**
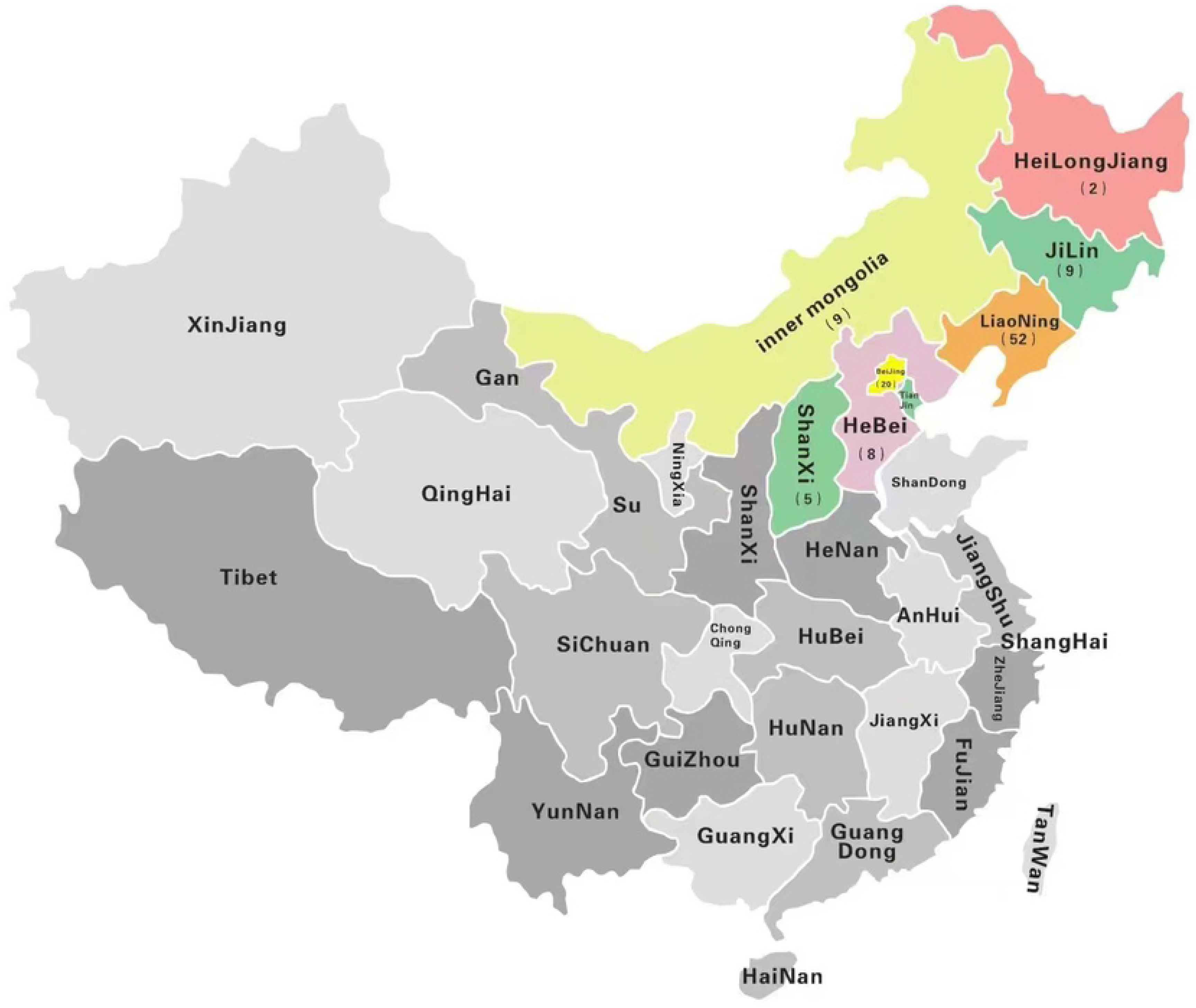
Geographic origin of millet (*Setaria italica* [L.]) germplasm.

### 2.2 Experimental design

The experiment was conducted over three growing seasons, from 2018 to 2020, at the experimental site of the Cash Crops Institute of Liaoning, located in Liaoyang City, Liaoning Province. The soil at the site is sandy loam, with the following composition in the topsoil: 1.97% organic matter, 0.08% total nitrogen, 73.4 mg/kg alkali-hydrolyzable nitrogen, 23.6 mg/kg available phosphorus, and 247.5 mg/kg available potassium. The resources were arranged sequentially during sowing, with each resource planted in four rows with 3~4 cm plant spacing The length of each row was 5 m, with a spacing of 50 cm, resulting in a plot area of 10 square meters. Sowing took place around May 11^th^, and harvesting was performed around September 25^th^. A three-compound fertilizer (N:P:K=15%:15%:15%) was applied as a base fertilizer. Ploughing and weeding were conducted three times during the growth period.

Two points were selected from each plot, and five consecutive plants with uniform growth were chosen from each point as samples. Quality and quantitative traits were investigated following the Descriptors and Data Standard for Millet (*S. italica* [L.]) compiled by Lu^[14]^. A total of 12 quantitative traits and 19 quality traits were selected for statistical analysis,ignoring traits that show no difference between varieties. The mean value of the ten plants and the interannual mean value were calculated.

### 2.3 Data processing

Data collection and analysis were performed using EXCEL 2007 and DPS 9.5^[15]^ Phenotypic diversity analysis and systematic cluster analysis(SCA)were conducted based on all 31 traits, while the Shannon diversity index was analysed using the method described by Wang^[16]^. Correlation analyses, principal component analyses(PCA), gray relational analysis(GRA), and K-means cluster analysis were performed based on the 12 quantitative traits.

Before conducting SCA and GRA, all the data were standardized using the following formula: 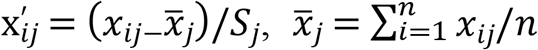, and 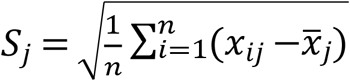. After standardization, a new data series was obtained with a dimension of 1, a mean value of 0 and a variance of 1.

## 3. Results and analysis

### 3.1 Phenotypic diversity analysis of millet germplasm resources

Table 1 shown the Shannon diversity index (DI) and distribution frequency of phenotypic characters of 19 quality traits. By comparing the frequency, it was found that two traits—tiller habit (TH) and branch habit (BH)—showed weak, moderate, and strong characteristics with a uniform distribution, resulting in a greater diversity index (DI). The phenotypic distribution of the other 17 traits showed clear tendencies, with a focus on 1 or 2 characteristics, resulting in a lower DI. It is worth noting that traits related to pigmentation show obvious distribution tendencies, with traits related to leaves mainly being green and traits related to flowers primarily being yellow.

**Table 1:**
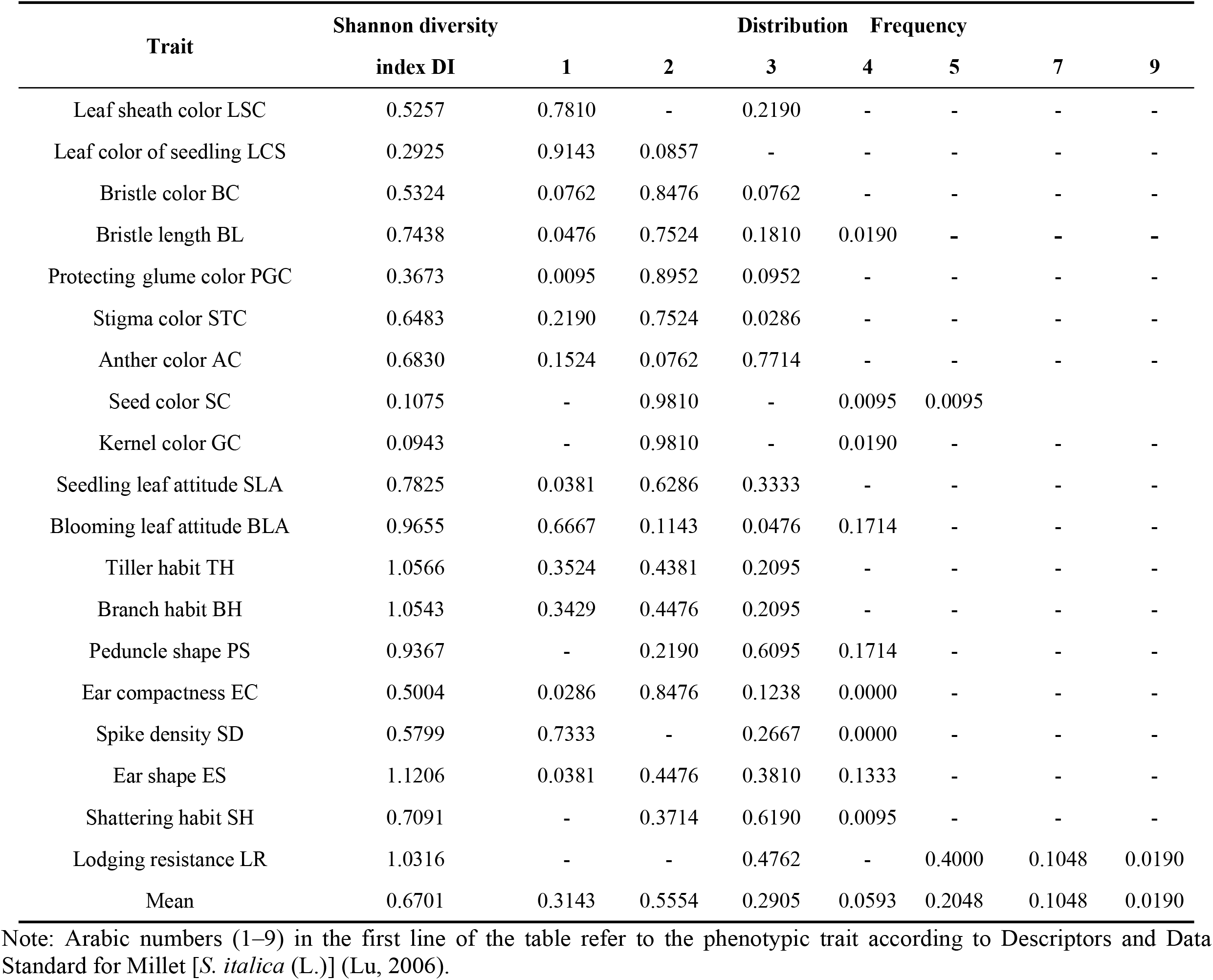
Phenotypic diversity of 19 quality traits.

The phenotypic diversity analysis of 12 quantitative traits is presented in Table 2. The coefficient of variation (CV), which represents the degree of trait dispersion, ranged from 6.34% to 43.77% for the 12 traits. The highest CV was observed for stem number per plant (SNP), followed by main stem node number (MSN), seed weight per plant (SWP), and ear weight per plant (EWP), indicating a high degree of variation and potential for genetic improvement. On the other hand, the subear internode length (SIL) and growing period (GP) had CVs close to or less than 10%, suggesting lower dispersion and relatively stable performance among the varieties. The Shannon DI, which reflects the distribution of trait performance, ranged from 1.0566 to 2.0428. The main stem length (MSL), EWP, SWP, and grass weight per plant (GWP) had DI values close to or greater than 2.0, indicating these traits performance are concentrated, reflecting a simple genetic basis,. Conversely, SNP had the lowest DI value, close to 1.0,indicating these traits performance were relatively scattered and susceptible to external conditions

**Table 2:** Phenotypic diversity of 12 quantitative traits.

|  | Traits | Min V. | Max V. | Mean V. | SD | CV % | Shannon diversity index DI |
| --- | --- | --- | --- | --- | --- | --- | --- |
|  | Growing period GP d | 101.0 | 131.0 | 118.809 5 | 7.535 9 | 6.34 | 1.696 6 |
| Stem | Stem number per plant SNP | 1.0 | 3.0 | 1.771 4 | 0.775 3 | 43.77 | 1.056 6 |
|  | Main Stem Length MSL cm | 90.0 | 170.0 | 133.847 6 | 19.025 2 | 14.21 | 2.042 8 |
|  | Main stem diameter MSD mm | 15.0 | 41.0 | 25.895 2 | 4.235 7 | 16.36 | 1.850 7 |
|  | Main stem nod number MSN | 0.3 | 1.0 | 0.522 4 | 0.150 6 | 28.83 | 1.517 6 |
| Ear | Peduncle length PL cm | 9.0 | 16.0 | 13.115 4 | 1.430 0 | 10.90 | 1.707 4 |
|  | Main ear length MEL mm | 16.0 | 40.0 | 22.847 6 | 4.018 7 | 17.59 | 1.874 1 |
|  | Main ear diameter MED mm | 0.7 | 4.2 | 2.635 2 | 0.503 6 | 19.11 | 1.764 9 |
| Yield | Grass weight per plant GWP g | 9.2 | 119.8 | 47.040 0 | 21.563 6 | 45.84 | 1.951 3 |
|  | 1000-seed weight TSW g | 2.0 | 3.9 | 2.688 6 | 0.350 1 | 13.02 | 1.578 4 |
|  | Ear weight per plant EWP g | 11.5 | 41.1 | 22.511 4 | 5.948 4 | 26.42 | 1.996 7 |
|  | Seed weight per plant SWP g | 9.3 | 27.3 | 15.514 3 | 4.104 3 | 26.45 | 1.957 0 |

### 3.3 Correlation analysis of millet germplasm resources

Correlation analysis aims to study the association between two or more random variables of equal status. In this experiment, correlation analysis was performed based on the 12 quantitative traits (Table 3). The EWP showed a significant positive correlation with SWP and both significant positive correlated with the main ear length (MEL), main ear diameter (MED), and SNP. also showed a highly significant negative correlation with the MSN. EWP had a significant positive correlation with MSL and GP, while SWP showed significant positive correlations with these traits. These results indicate that grain yield is positively correlated with ear size (MEL and MED) and ear setting potential (SNP and GP). Additionally, GWP was positively correlated with the MSN and main stem diameter (MSD), which represent vegetative growth. GP, which represents the growth potential of the plant, showed positive correlations with SNP, MSL, MEL, SIL, and MED. Overall, the results of the correlation analysis align with expectations.

**Table 3:** Correlation analysis of 12 quantitative traits.

| Correlation Coefficient | SNP | MSL | PL | MSD | MSN | MEL | MED | GP | GWP | TSW | EWP |
| --- | --- | --- | --- | --- | --- | --- | --- | --- | --- | --- | --- |
| SNP | 1 |  |  |  |  |  |  |  |  |  |  |
| MSL cm | 0.023 7 | 1 |  |  |  |  |  |  |  |  |  |
| PL cm | 0.083 4 | 0.428 4** | 1 |  |  |  |  |  |  |  |  |
| MSD cm | 0.196 5 | -0.150 6 | 0.082 8 | 1 |  |  |  |  |  |  |  |
| MSN | -0.169 3 | 0.088 1 | 0.017 7 | -0.075 9 | 1 |  |  |  |  |  |  |
| MEL mm | 0.223 3* | 0.325 5** | 0.194 5* | 0.246 3* | -0.215 1 | 1 |  |  |  |  |  |
| MED mm | 0.262 2** | -0.008 1 | -0.007 3 | 0.180 9 | -0.093 5 | 0.005 1 | 1 |  |  |  |  |
| GP d | 0.410 5** | 0.302 8** | 0.199 7* | 0.097 8 | -0.095 6 | 0.400 0** | 0.192 0* | 1 |  |  |  |
| GWP g | -0.010 7 | 0.081 1 | 0.075 9 | -0.194 5* | 0.287 2** | 0.042 7 | -0.105 8 | 0.055 8 | 1 |  |  |
| TSW g | -0.190 4 | -0.044 6 | -0.027 4 | -0.017 0 | 0.052 5 | -0.028 6 | -0.045 1 | -0.024 2 | -0.050 8 | 1 |  |
| EWP g | 0.395 5** | 0.292 7** | 0.182 1 | 0.093 1 | -0.284 6** | 0.461 2** | 0.279 9** | 0.330 3** | -0.160 9 | -0.095 5 | 1 |
| SWP g | 0.253 3** | 0.214 7 * | 0.177 7 | 0.003 5 | -0.267 2** | 0.376 2** | 0.257 3** | 0.193 8* | -0.092 1 | -0.083 9 | 0.826 6** |

Note: $r_{0.05}=0.1918$ , $r_{0.01}=0.2504$

### 3.4 Principal component analysis (PCA) of quantitative traits

Principal component analysis (PCA) is a comprehensive method used for transforming multiple trait indices and reducing dimensionality into several principal components. Each principal component represents a relatively independent indicator system, there is no correlation between each principal component, the numerical value is intuitive and easy to analyze^[17]^. The PCA results for 12 quantitative traits from 105 millet germplasms are presented in Table 4. Seven principal components with eigenvalues greater than or close to 1 were selected, which collectively contributed to 81.63% of the variance and encapsulated most of the genetic information of the millet germplasms^[18-19]^ (Qiao, 2015; Liu, 2020).

**Table 4:**
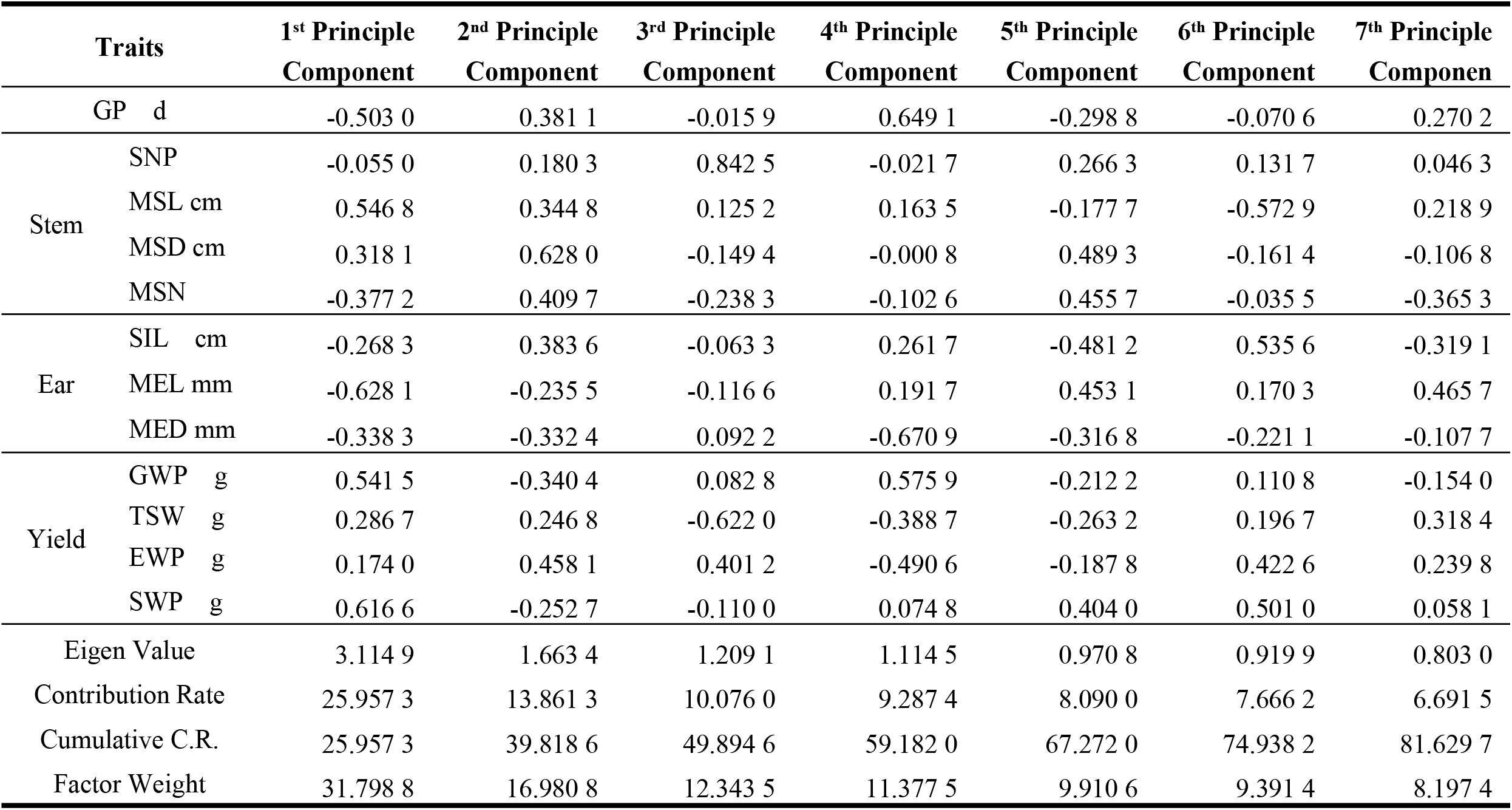
Principal components and eigenvalues of 12 quantitative traits.

**Table 5:**
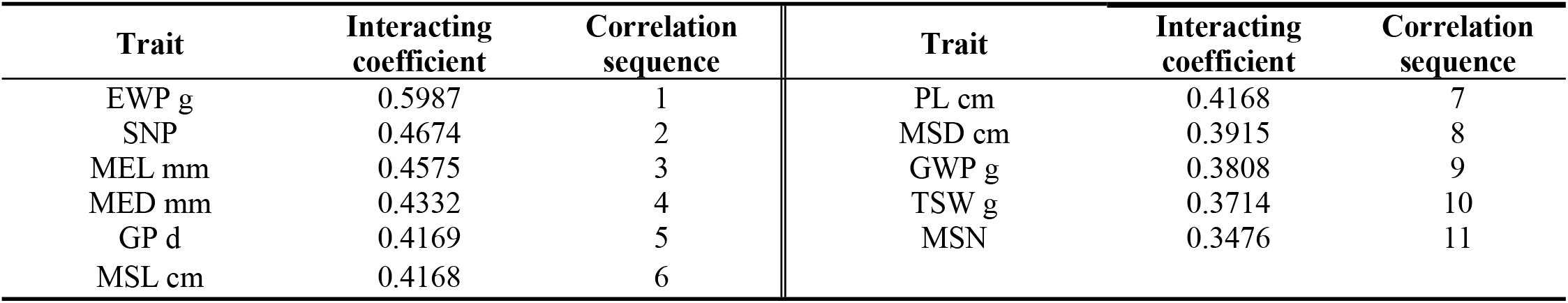
Gray relational analysis of seed yield and 11 quantitative traits.

| Trait | Interacting coefficient | Correlation sequence | Trait | Interacting coefficient | Correlation sequence |
| --- | --- | --- | --- | --- | --- |
| EWP g | 0.5987 | 1 | PL cm | 0.4168 | 7 |
| SNP | 0.4674 | 2 | MSD cm | 0.3915 | 8 |
| MEL mm | 0.4575 | 3 | GWP g | 0.3808 | 9 |
| MED mm | 0.4332 | 4 | TSW g | 0.3714 | 10 |
| GP d | 0.4169 | 5 | MSN | 0.3476 | 11 |
| MSL cm | 0.4168 | 6 |  |  |  |

The eigenvector value of each trait indicates its contribution to the principal component (PC). By comparing the eigenvector values, the following observations were made: the 1^st^ PC was primarily associated with MEL(-)-, MSL(+)-, seed weight per plant (SWP +), GWP(-) -, and GP(-) -; the 2^nd^ PC was mainly influenced by MED(+), EWP(+), and MSN(+); the 3^rd^ PC was primarily influenced by SNP(+), 1000-seed weight (TSW-), and EWP(+). the 4^th PC^ was mainly influenced by MED(-), GP(+), GWP(+) and EWP(-); the 5^th^ PC was mainly influenced by MSD(+), SIL(-), MSN (+), MEL (+) and SWP(+); the 6^th^ PC was primarily influenced by MSL (-), SIL (+), SWP (+), and EWP (+); and the 7^th^ PC was mainly influenced by MEL (+) and MSN (-). Compared to correlation analysis, PCA provided an interpretation of the correlation between quantitative traits of plants and specifically emphasized the importance of the GP and TSW.

### 3.5 Gray relational analysis (GRA) of quantitative traits and yield

Gray correlation analysis refers to the quantitative description and comparison of the development of a system to measure the degree of correlation between factors based on the similarities or differences in their development trends^[20]^. After standardized processing, the seed weight per plant (SWP) was taken as the reference column, and the correlation between the main quantitative traits and the SWP was analysed. Among the tested quantitative traits, EWP had the greatest impact on the SWP, followed by SNP, MEL, and MED, these four traits directly represent the ear-bearing capacity of the plant. The coefficients of three traits, namely, GP, MSL, and SIL, were very close, these three traits are mainly related to maturity and main stem growth. The MSD, GWP, and MSN represent the vegetative growth status of the plant, and their contributions to the SWP decrease in turn. Therefore, when selecting high-yield millet varieties, it is advisable to focus on materials with more tillers, larger ears, and longer growth periods.

Cluster analysis reflects the genetic differences among different varieties, and clustering resources with similar traits into one group can guide the selection of hybridization parents^[21]^. The selection of resources from different groups with large genetic distances for hybridization can lead to substantial genetic variation^[22-23]^.

Based on the performance of all 31 traits (12 quantitative traits + 19 quality traits) of 105 millet resources, SCA analysis was conducted via standardized data transformation - Euclidean distance - deviation square sum method. All materials were divided into six groups at Euclidean distance = 17.09 (Fig. 2).

**Fig. 2:**
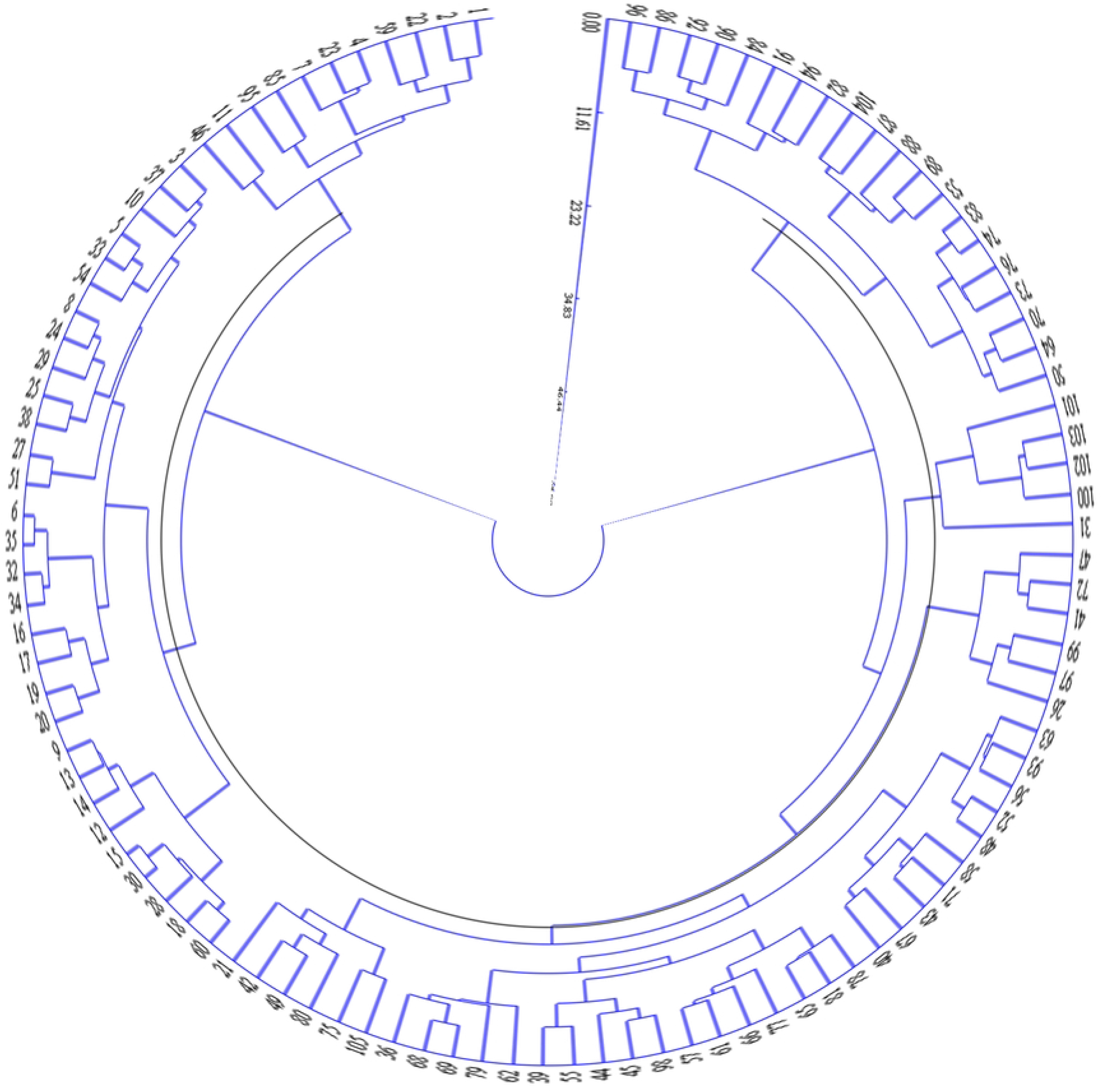
System cluster analysis of Setaria italica (L.) germplasm.

The quantitative traits of different groups were analysed and compared. The results are presented in Table 6. The 1^st^ group consisted of 11 samples, which exhibited mid-earlier maturity, fewer tillers, taller and slender main stem with multiple nodes, smaller main ears, small seeds with medium grain yield, and higher grass yield. The 2^nd^ group included 30 materials, showing earlier maturation, fewer tillers, lower and slenderer main stem with multiple nodes, smaller main ears, larger seeds but lower grain yield, and higher grass yield. The 3^rd^ group consisted of 33 materials displaying mid-later maturation, medium tiller ability, taller main stem with multiple nodes, medium-sized main ears, larger seeds with higher grain yield, and medium grass yield. The 4^th^ group comprised 6 materials, showing later maturation, multiple tillering, taller and stronger main stems with multiple nodes, larger main ears, smaller seeds but higher grain yield, and greater grass yield. The 5^th^ group contained 5 materials, exhibiting mid-later maturation, multiple tillering, lower and stronger main stem with fewer nodes, shorter and thicker main ears, medium-sized grains, lower grain yield and lower grass yield. The 6^th^ group consisted of 20 materials that exhibited mid-later maturation, multiple tillering, taller and slenderer main stems, smaller main ears, smaller seeds but higher grain yields, and lower grass yields.

**Table 6:**
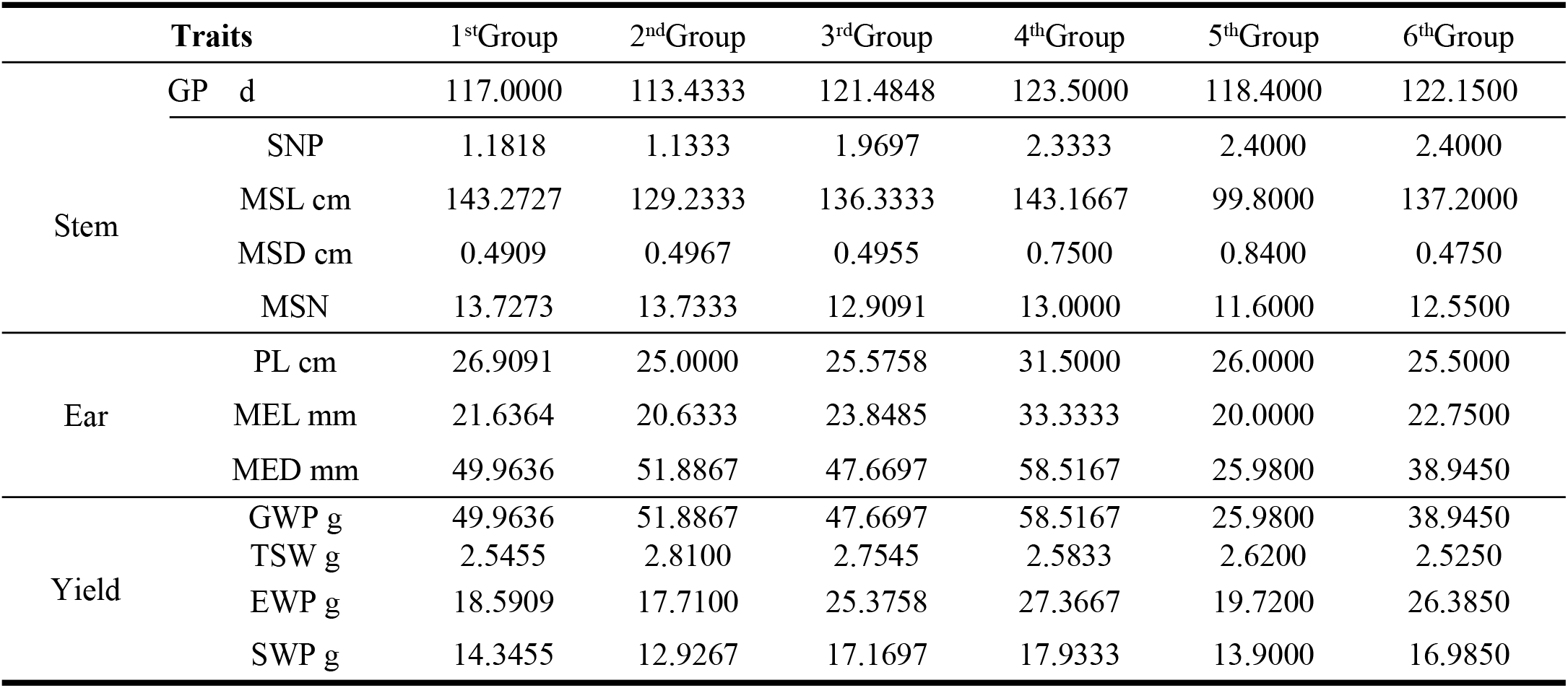
System cluster analysis of 12 quantitative traits.

In the analysis of 12 quantitative traits in Table 6, the qualities of each group were also compared. The 1^st^ group had unique characteristics in terms of leaf sheath color (LCS) and bristle length (BL) that were not found in the other groups. The 2^nd^ group displayed common characteristics for all traits. The 3^rd^ group exhibited abundant traits characteristics, including red seed color (SC), which was not present in the other groups. The 4^th^ group, with its 6 resources, had relatively simple and concentrated trait phenotypes but also had a rare occurrence of purple bristle color (BC). The 5^th^ group, consisting of only 5 resources, had scattered trait phenotypes, with kernel color (KC), SC, and SH showing unique characteristics. The 6^th^ group showed abundant trait characteristics, with LR and ear shape (ES) displaying characteristics not found in the other groups.

Further examination of the geographical distribution of resources within each group revealed that resources from Liaoning had certain advantages, particularly in Group 4, which exclusively consisted of materials from Liaoning. Resources from Group 1 were also concentrated in Liaoning. Group 2 resources were relatively dispersed and covered almost all geographical origins. Groups 3 and 6 mainly sourced their resources from Liaoning, Beijing, and Inner Mongolia. Group 5 resources were evenly distributed from Liaoning and Jilin.

### 3.6 K-mean clustering analysis

K-means clustering analysis was conducted to analyse the 12 quantitative traits, and the theoretical distribution intervals and central values of these traits are shown in Table 7. The performance of each trait for the studied resources was found to be concentrated near the central value, with more distribution around the minimum value and less around the maximum value.

**Table 7:**
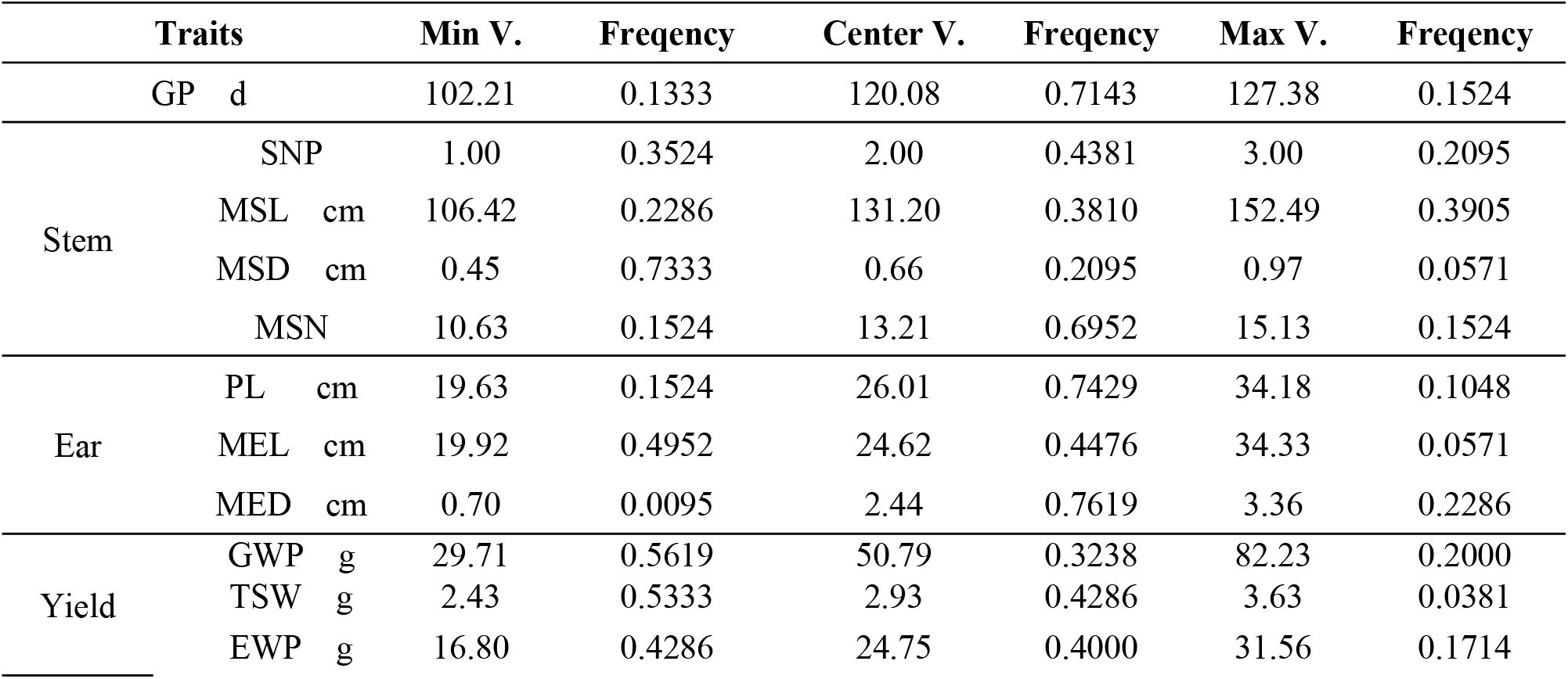
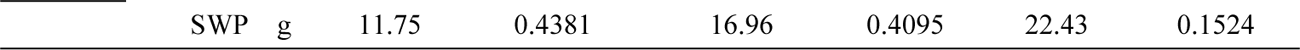
K-mean cluster analysis of 12 quantitative traits.

Based on these results, specific resource materials with desirable performance in terms of maturity, plant height, tiller habit, ear size, and grain weight were identified. After a comprehensive evaluation, four specific resources were selected: Dungu No. 1, an early-maturing and draft small-ear and small-seed resource from Taiyuan, Shanxi Province; Xiao-li-xiang, a small-seed and early-maturing resource from Shijiazhuang, Hebei Province; Basen Shengu, a later-maturing and taller large-ear resource from Fuxin, Liaoning Province; and Yuhuanggu No. 1, a later-maturing and low-yield large-seed resource from Chifeng, Inner Mongolia.

## 4. DISCUSSION

This experiment examined a total of 31 traits, which showed rich diversity in both qualitative and quantitative aspects. Among the 19 qualitative traits, traits related to plant color, such as KC, SC, and LSC, exhibited lower Shannon DI values, suggesting a noticeable tendency toward pigmentation in the local resources. Leaf-related traits are shown in green, while ear-related traits are shown in yellow. Traits representing ear characteristics, such as ear compactness (EC, SD), bristle length (BL), displayed moderate DI values, indicating that the ears of local resources are primarily sparse and loose, making them prone to grain drop. The BL was short, and the peduncle shape was curved, with the ear shape predominantly cylindrical and spindle shaped. Traits representing plant structure, such as TH, BH, and blooming leaf attitude (BLA), showed higher DI values, indicating greater diversity in plant structure types.

For the 12 quantitative traits examined, the DI values ranged from 1.0566 to 2.0428, and the CVs ranged from 6.34% to 45.84%. GP and TSW exhibited small DI values, suggesting strong adaptability through long-term selection and limited potential for genetic improvement. This finding is consistent with the results obtained by Gao^[24]^ in their study on mung beans. Yield traits, such as EWP, SWP, and GWP, which reflect plant growth capacity, exhibited relatively high CV and DI values. This indicates that these traits are influenced by multiple quantitative genes, are susceptible to external conditions, and have great potential for genetic improvement. This conclusion aligns with the research findings^[25-26]^. The results of the correlation analysis indicate that ear size, SNP, and GP are closely correlated with grain yield, while no direct correlation is found between traits representing plant growth status and grain yield. These findings are consistent with those of Jia^[27]^ but differ from those of Yan^[28]^, which can be attributed to variations in test sites, sampling methods, and measurement indices. Additionally, the correlation of qualitative traits was also analyzed by assigning values based on their phenotype (result supplied), revealing the correlation between traits associated with pigmentation, and represented BH, TH, Spike density (SD), Shattering habit (SH), Seedling leaf attitude (SC) and Lodging resistance (LR) exhibited correlations with yield, consistent with the results of the diversity analysis.Since the correlation of qualitative traits is completed by assigning values on each phenotype, human factors influence greatly, the analysis results can be provided as a reference in the work only.

PCA revealed the extraction of 7 PCs. The 1st, 2nd, 3rd, 4th, and 6th PCs focus on yield and explain the associations between yield and main stem growth ability, main ear length, and growth period. The 5th and 7th PCs highlight the relationship between main stem growth ability and main ear length. In the breeding process, it is important to consider the contribution rate of each PC and the breeding goal comprehensively.

GRA based on GWP demonstrated that ear size and ear-bearing capacity are the main factors limiting yield, followed by growth period traits.

Overall, the results of correlation analysis, PCA, and GRA were consistent, suggesting that high-yield lines are expected to have more tillers and branches, larger and well-formed ears, and lodging resistance. It is worth noting that TSW does not directly influence yield, possibly due to millet’s seed shatter characteristics and the presence of immature seeds at harvest. These findings are similar to those obtained by Jia^[29]^ in their study on Tartary buckwheat, where the length of subear internode (SIL) was found to be proportional to the GP, making it a potential indicator of maturity.

A systematic cluster analysis was conducted on all 31 traits, resulting in the division of 105 millet materials into six groups at a Euclidean distance of 17.09. Each group exhibited distinct phenotypic characteristics and showed certain geographical distribution tendencies. According to previous studies on crops such as adzpea^[30]^ and cotton^[31]^ (Xu et al., 2017), the characteristics of these groups were found to be correlated with the natural climate conditions of their original source areas, which could be categorized using cluster analysis. The geographical distribution tendencies among groups were not prominently observed in this experiment, due to the frequent introduction of resources between regions and intermixing of bloodlines.

In addition, K-means cluster analysis was performed on 12 quantitative traits. Theoretical distribution intervals and central values were calculated to identify resource materials with specific traits. Fuether more,, four specific resources were identified: one with early maturity and draft, small ear size, and small seeds; one with small seeds and early maturity; one with later maturity, taller and larger ears; and one with later maturity,,lower yields, and larger seeds.

## 5. Conclusion

In conclusion, this study evaluated the phenotypic diversity of 105 millet germplasms in the Liaoning area by considering 19 qualitative traits and 12 quantitative traits. The results revealed a rich diversity of traits and complex correlations among them. Resources that possess characteristics such as upthrow seedling leaves, more tillers and branches, larger and well-formed ears, and lodging resistance prefer to higher grain yield. It was also discovered that the subear internode length (SIL) could be an indicator for maturity selection. Furthermore, all resource materials were divided into six groups with different phenotypic characteristics, and the distribution interval of each quantitative character was determined. Four specific resources, namely, Dungu No. 1, Xiao-li-xiang, Basen Shengu, and Yuhuanggu No. 1, were identified for further breeding and practical applications.

## Author Contribution Statement

LI X. T., HE W.F., WANG H.H. performed experiments.

HE W.F. supplied germplasm resources materials.

WANG H.H. analyzed data.

LI X.T. wrote the paper.

XU M. planned and supervised the research.

## Funding

This study was financially supported by philosophy and Social Sciences Planning Project Guizhou (Grant number:21GZYB62)

## Conflict interest

There are no direct competition and conflict interest of our work.

## Notes

### Competing Interest Statement

The authors have declared no competing interest.

